# Kidney Function and Blood Pressure: A Bi-directional Mendelian Randomisation Study

**DOI:** 10.1101/856674

**Authors:** Zhi Yu, Josef Coresh, Guanghao Qi, Morgan Grams, Eric Boerwinkle, Harold Snieder, Alexander Teumer, Cristian Pattaro, Anna Köttgen, Nilanjan Chatterjee, Adrienne Tin

**Author notes:** **Corresponding Author:** Adrienne Tin, PhD, Johns Hopkins Bloomberg School of Public Health, 615 N. Wolfe Street, Room E6608, Baltimore, MD 21205.

## Abstract

**Objective:** To evaluate the bi-directional causal relation between kidney function and blood pressure.

**Design:** Mendelian randomisation study.

**Setting:** We performed two-sample Mendelian randomisation analyses. Genetic instruments of kidney function traits were selected from summary statistics of genome-wide association studies (GWAS) of glomerular filtration rate estimated from serum creatinine (eGFRcr) and blood urea nitrogen (BUN) and were required to be associated with both eGFRcr and BUN to ensure that the instruments were more likely to represent the underlying kidney function. Genetic instruments of blood pressure were selected from summary statistics of GWAS of systolic and diastolic blood pressure. We investigated Mendelian randomisation hypothesis using several alternative approaches, including methods that are most robust to the presence of horizontal pleiotropy.

**Participants:** The summary statistics of eGFRcr included 567,460 participants from 54 cohorts, and the summary statistics of BUN included 243,031 participants from 48 cohorts from the Chronic Kidney Disease Genetics (CKDGen) Consortium. The summary statistics of systolic and diastolic blood pressure included 757,601 participants from the UK Biobank and 78 cohorts from the International Consortium for Blood Pressure (ICBP).

**Results:** Significant evidence supported the causal effects of higher kidney function on lower blood pressure with multiple methods. Based on the mode-based Mendelian randomisation analysis approach, known for its robustness to the presence of pleiotropic effect, the effect estimate for 1 SD higher in eGFRcr was −0.17 SD unit (95 % CI: −0.09 to −0.24) in systolic blood pressure (SBP) and −0.15 SD unit (95% CI: −0.07 to −0.22) in diastolic blood pressure (DBP). In contrast, the causal effects of blood pressure on kidney function were not statistically significant.

**Conclusions:** Mendelian randomisation analyses support causal effects of higher kidney function on lower blood pressure. These results suggest preventing kidney function decline can reduce the public health burden of hypertension.

## INTRODUCTION

Hypertension and chronic kidney disease (CKD) are two interconnected global public health burdens. The estimated prevalence of hypertension is as high as 31%, while CKD affects ∼10% of adults^1−3^. Both CKD and hypertension are major risk factors for cardiovascular disease (CVD) and mortality^4−6^. Hypertension has long been considered as a risk factor for kidney function decline and the development of CKD based on observational studies^7−10^. Reports on the association between kidney function decline and incident hypertension have been more limited^11 12^. Analysis of the causal effects of lower kidney function on higher blood pressure has inconsistent results^13^. Evaluating the causal relations between kidney function and blood pressure can inform disease prevention and treatment strategies.

Mendelian randomisation is an approach employing genetic variants as instrumental variables of the exposure to estimate causal effects between an exposure and an outcome with the goal of overcoming the confounding inherent in observational studies^14^. Using a Mendelian randomisation analysis approach, Liu *et al.* found that higher genetically-predicted systolic blood pressure (SBP) is causally linked to CKD^15^. Haas *et al.* showed evidence supporting the existence of a feed-forward loop between albuminuria, a marker of kidney damage, and hypertension^16^. Morris et al. reported significant causal effect of lower kidney function on higher diastolic blood pressure (DBP) and not on systolic blood pressure (SBP)^13^. Two-sample Mendelian randomisation analysis is an extension of the Mendelian randomisation method that allows the use of summary statistics of genome-wide association studies (GWAS) for conducting Mendelian randomisation studies. We performed two-sample bidirectional Mendelian randomisation analyses to assess the causal effects of kidney function on blood pressure and vice versa using summary statistics from large-scale GWAS. The primary kidney function trait was estimated glomerular filtration rate based on serum creatinine (eGFRcr)^17^. The primary blood pressure trait was SBP with DBP as secondary.

To obtain robust conclusions from our analyses, we paid particular attention to two critical aspects in this Mendelian randomisation study. One being the use of serum creatinine for GFR estimation, which might link eGFRcr to genetic variants more related to creatinine metabolism than glomerular filtration function, making it difficult to interpret any causal findings between eGFRcr and blood pressure. To address this issue, we used additional data from large-scale meta-analysis of GWAS of blood urea nitrogen (BUN), an alternative kidney function biomarker, to select genetic instruments that are likely more specific to kidney function. The second being the assumption of the lack of horizontal pleiotropy of the genetic instruments variants, which is usually difficult to assess and verify^18^. To address this issue, we analyzed the data using multiple Mendelian randomisation methods and prioritized the method that are known to be most robust to the presence of horizontal pleiotropy^19^.

## METHODS

### Study design overview

Two-sample Mendelian randomisation allows for the estimation of causal effects using GWAS summary statistics of the exposure and the outcome from different populations without using individual level data. We performed two-sample Mendelian randomisation analyses to estimate the causal effects of kidney function on blood pressure and vice versa. The primary kidney function trait was eGFRcr with BUN as a secondary trait. CKD, defined as eGFRcr < 60 mL/min/1.73m^2^, was a secondary outcome^20^. The primary blood pressure trait was SBP with DBP as secondary^21^. Published GWAS summary statistics were obtained from European-ancestry participants of the Chronic Kidney Disease Genetics (CKDGen) Consortium^20^ for kidney function and the UK Biobank and International Consortium for Blood Pressure (UKB-ICBP)^22^ for blood pressure. All GWAS summary statistics assumed an additive genetic model.

### Summary statistics of kidney function from the CKDGen Consortium

The meta-analysis of the GWAS of eGFRcr included 54 cohorts of European ancestry (N = 567,460), largely adult population-based (median age among the cohorts: 53.4 year; median of % male: 48%). A small proportion of the participants were from cohorts of CKD patients, diabetes patients, or children (2.5%). The meta-analysis of the GWAS of BUN included 48 cohorts of European ancestry (N = 243,031), and the analysis of CKD included 444,971 participants. eGFRcr was calculated using the Chronic Kidney Disease Epidemiology Collaboration (CKD-EPI) equation^17^ for adults and the Schwartz formula^23^ for participants who were 18 years old or younger. BUN, the secondary kidney function trait, was derived as blood urea×2.8, with units expressed as mg/dl^20^. The phenotypes used in the GWAS of eGFRcr and BUN were the natural log transformed age- and sex-adjusted residuals of the traits. Genotypes were imputed using the Haplotype Reference Consortium (HRC)^24^ or the 1000 Genomes Project^25^ reference panels.

### Summary statistics of blood pressure from the UKB-ICBP

Summary statistics of blood pressure traits were obtained from the combined meta-analysis results of the UK Biobank (UKB) and the International Consortium of Blood Pressure Genome Wide Association Studies (ICBP)^22^. The UKB is a population-based cohort with ∼500,000 participants (mean age: 57; female: 54%) with deep genetic and phenotypic data, including blood pressure measurements^26^. SBP and DBP were calculated as the mean of two automated or manual blood pressure measurements, except for a small number of participants with one blood pressure measurement (n = 413). The GWAS of SBP and DBP in UKB included 458,577 participants of European ancestry. Genotypes were imputed using the HRC reference panel^22 24^. The meta-analysis of the GWAS of SBP and DBP from ICBP included 77 cohorts of European ancestry (N = 299,024)^21^. Genotypes were imputed using the HRC^24^ or the 1000 Genomes Project^25^ reference panels^22^. In both UKB and ICBP, the values of SBP and DBP were adjusted for the use of blood pressure lowering medications by adding 15 and 10 mmHg, respectively^22 27^.

### Mendelian randomisation assumptions

Genetic instruments used in Mendelian randomisation studies rely on three assumptions: (i) the SNP must be associated with the exposure; (ii) the SNP is independent of confounders, i.e. other factors that can affect the exposure-outcome relationship; and (iii) the SNP must be associated with the outcome through the exposure only, i.e., no direct association due to horizontal pleiotropy^28^.

### Selection of genetics instruments more likely to be related to kidney function

To ensure that the genetic instruments satisfied the first assumption with respect to kidney function, we selected index SNPs associated with multiple biomarkers of kidney function so that they are more likely to be related to GFR, the exposure of interest, rather than the GFR biomarker. For the primary analysis using eGFRcr, we started with the index SNPs of the genome-wide significant loci of the European-ancestry meta-analysis of eGFRcr from the CKDGen Consortium^20^. We first evaluated the association of the index SNPs with potential confounders using the GWAS summary statistics from UKB for the following traits: prevalent diabetes, body mass index (BMI), triglycerides and high-density lipoprotein cholesterol (HDL-C) levels, smoking, and prevalent coronary heart disease^1 29 30^. We removed index SNPs with genome-wide significant associations (5×10^−8^) in UKB with the potential confounders listed above^1 29 31^

We then used genetic association information of BUN^20^, an alternative biomarker of kidney function, to select genetic instruments that were more likely to reflect kidney function as opposed to creatinine metabolism. This approach was similar to the approach in Wuttke *et al.* for prioritizing genetic loci most likely to be relevant for kidney function^20^. We required that the index SNPs selected from eGFRcr GWAS to be associated with BUN at a Bonferroni-corrected significance (p < 0.05 divided by the number of eGFRcr index SNPs) and in opposite direction since higher GFR would lead to lower BUN. To ensure independence among genetic instruments, we applied pairwise-linkage disequilibrium (LD) clumping^32^ with a clumping window of 10 MB and an *r*^2^ cutoff of 0.001 (default of the clump_data function)^32^. The matching of the effect allele of each SNP between the summary statistics of the exposure and the outcome was examined using the harmonise_data function, which removed SNPs that were palindromic or had possible strand mismatch. Finally, to reduce the possibility that a genetic instrument might affect the outcome independently of the exposure, we applied Steiger filtering to ensure that the association between a genetic instrument and the exposure was stronger than its association with the outcome^33^.

To select genetic instruments of BUN, the secondary kidney function trait, we started with index SNPs identified from GWAS of BUN and followed similar procedure of selection. We used their association with eGFRcr for screening out those SNPs that might only be related to metabolism of BUN but not to kidney function.

### Selection of genetics instruments for blood pressure

For blood pressure traits, we started with the index SNPs from genome-wide significant loci of SBP or DBP reported by the UKB-ICBP^22^, applied the same steps as described above for eGFRcr, without the alternative biomarker step. Briefly, we removed index SNPs that were associated with potential confounders listed above, removed correlated SNPs using the clump_data function^32^, used the harmonise_data function to remove SNPs that were palindromic or had possible strand mismatch between the summary statistics of the exposure and outcome, and finally, we applied Steiger filtering^33^

### Use of robust method to account for horizontal pleiotropy

It is well known a number of existing methods for Mendelian randomisation analysis can be heavily biased in the presence of direct association of SNP with the outcome that is not mediated by the exposure^34^. In particular, when the direct effects of genetic instrument on the outcomes and the exposures are correlated across different instruments due to the presence of unobserved confounders that may have heritable components, the bias can be severe^19^. Thus, to reduce the possibility that the genetic instruments might affect the outcome independently of the exposure, in addition to the use of Steiger filtering^33^ discussed above, we chose the weighted mode method, known to be most robust in the presence of horizontal pleiotropy, as our primary Mendelian randomisation method. In addition, we conducted sensitivity analysis using a number of alternative methods that may be more powerful under various model assumptions (see Sensitivity Analysis section). Given our primary traits were eGFRcr for kidney function and SBP for blood pressure, the significance level for Mendelian randomisation analysis was set at p-value < 0.025.

### Units of causal effect estimates

For continuous exposures and outcomes, we estimated the causal effects of 1 standard deviation (SD) difference in the outcome per 1 SD higher in exposure. The effect estimates of the genetic instruments for the exposure and the outcome were scaled using the estimated SD of the trait. To be consistent with the outcome used in the GWAS of CKDGen, the SD of log(eGFRcr) and log(BUN) were estimated using the natural log transformed sex- and age-adjusted residuals among European-ancestry participants of the Atherosclerosis Risk in Communities Study (ARIC), a population-based study (n = 11,478, SD of log[eGFRcr residuals]: 0.13, SD of log[BUN residuals]: 0.24). The SD of SBP and DBP were estimated based on 474,382 participants of UKB (SD of SBP: 19.3 mmHg, SD of DBP: 10.5 mmHg)^35 36^.

### Sensitivity analyses

Several sensitivity analyses were used to evaluate the robustness of the causal effect estimates of kidney function on blood pressure and vice versa. First, in addition to weighted mode, our primary method, we estimated causal effects using alternative Mendelian randomisation methods: inverse-variance-weighted fixed-effects (IVW-FE) method^37^, Mendelian randomisation-Egger (MR-Egger)^38^, weighted median^39^, and Mendelian randomisation analysis using mixture models (MRMix)^40^, a novel method that uses a mixture model with components for valid and invalid instruments and then conducts a grid search to obtain the optimal estimate^40^. Second, given that CKDGen and ICBP have overlapping samples, which could potentially bias the causal effect estimates towards the observational effect^41^, we examined the bi-directional causal estimates between kidney function and blood pressure using GWAS summary statistics of blood pressure from UKB only, which was not part of the CKDGen meta-analysis of kidney function traits. Third, eGFR estimated from cystatin C (eGFRcys) is another alternative measure of kidney function with published GWAS results (n = 12,266)^42^. We evaluated whether using eGFRcys instead of BUN as the alternative biomarker would lead to similar results. For eGFRcr, we required that the genetic instruments of eGFRcr to be associated with eGFRcys in the same direction with Bonferroni-corrected significance. For BUN, we required the genetic instruments of BUN to be associated with eGFRcys in the opposite direction with Bonferroni-corrected significance. Power analysis were calculated by an online tool tailored for Mendelian randomisation (https://sb452.shinyapps.io/power/)^43^. All analyses were conducted using R (version 3.5.3), and the “TwoSampleMR” package was used for all Mendelian randomisation analyses, except MRMix.

## RESULTS

### Selection of kidney function genetic instruments

Of 256 reported eGFRcr index SNPs, 43 were removed due to association with potential confounders (**Figure 1, Supplementary Table 1**). Of the remaining 213 index SNPs, 40 satisfied our selection criteria using BUN as the alternative kidney function biomarker (**Supplementary Table 2**). For example, the index SNP at *GATM*, an enzyme in creatine metabolism^44^, was removed due to insignificant association with BUN (rs1145077, eGFRcr p = 6.9×10^−142^, BUN p = 0.92). After LD clumping and matching of coding and non-coding alleles between exposure and outcome, 35 index SNPs remained. Finally, Steiger filtering removed the index SNPs at *FGF5* and *SPI1* (**Supplementary Table 3**) resulting in 33 genetic instruments for eGFRcr. Using similar procedures, the number of genetic instruments retained for BUN was 24. For example, using eGFRcr as the alternative kidney function biomarker, the BUN index SNP at *SLC14A2*, a urea transporter^45 46^ was removed due to insignificant association with eGFRcr (rs41301139, p = 0.14) (**Supplementary Table 4)**. The numbers of SNPs retained after each selection step are reported in **Supplementary Table 5**.

**Figure 1.**
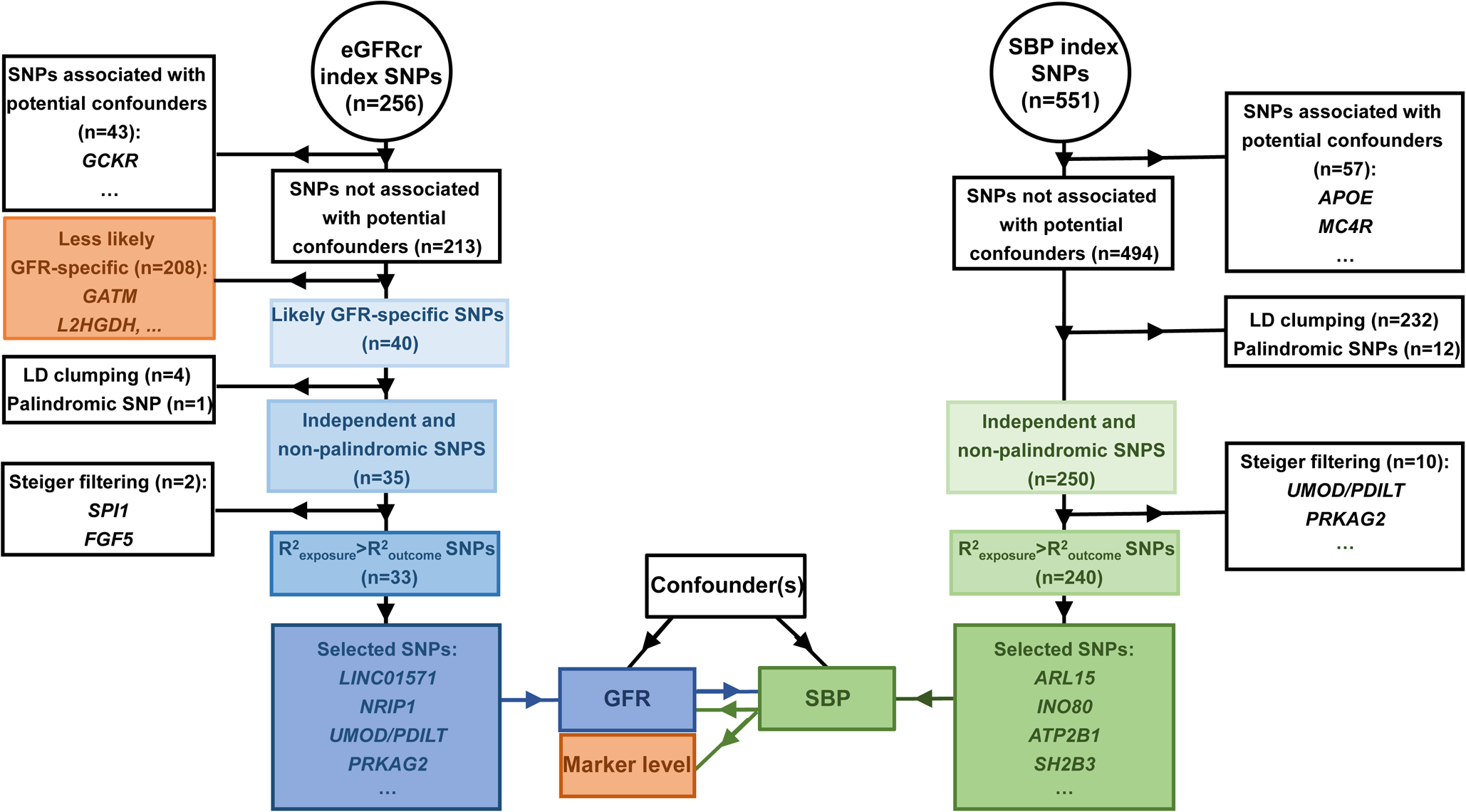
Selection of genetic instruments for eGFRcr and SBP. Details of the selection of genetic instruments are reported in Supplementary Table 1 (association with confounders), Supplementary Table 2 (use of BUN as alternative kidney function biomarker), Supplementary Table 3 (Steiger filtering), Supplementary Table 5 (summary of the number of index SNPs retained at each step).

### Significant causal effect of kidney function on blood pressure

We identified significant evidence for causal effects of higher kidney function for lower blood pressure. Using weighted mode, the primary method, the causal estimates for each SD higher log(eGFRcr) were −0.17 SD in SBP (95% confidence interval [CI]: −0.24 to −0.09; p = 9.92×10^−5^) and −0.15 SD in DBP (95% CI: −0.22 to −0.07; p = 5.02×10^−4^, **Figure 2**). These causal effects were equivalent to a 50% lower in eGFRcr leading to 17.5 mmHg higher SBP and 8.4 mmHg higher DBP. We also observed significant causal effects of BUN, the secondary kidney function trait, to SBP and DBP using the weighted mode method (SBP p = 4.92×10^−4^; DBP p = 3.88×10^−6^). Using other Mendelian randomisation methods, IVW-FE, MR-Egger, weighted median, and MRMix, all causal effect estimates were in the same direction as those from weighted mode and significant, providing support for causal effects of lower eGFRcr on higher SBP and DBP (**Supplementary Table 6**). The scatter plots with the regression line from all Mendelian randomisation methods are presented in **Supplementary Figures 1a to 4a.** The forest plots of single SNP effects from each of the kidney function traits to each of the blood pressure traits are presented in **Supplementary Figures 1b to 4b**.

**Figure 2.**
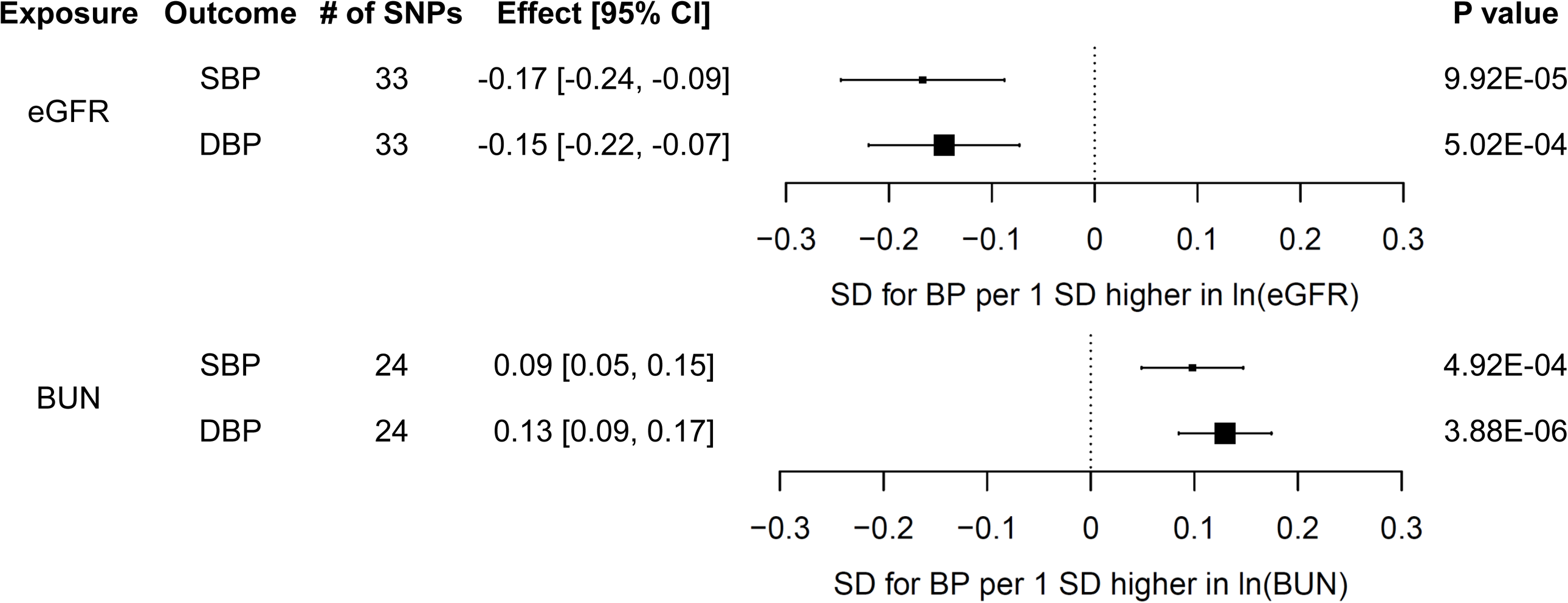
Estimates of the causal effects [95% confidence intervals] from eGFRcr on SBP and DBP (A) and BUN on SBP and DBP (B) using the weighted mode method

Sensitivity analysis using blood pressure summary statistics from UKB only as the outcome, without cohorts overlapped with CKDGen, resulted in similar significant causal estimates for eGFRcr and BUN on SBP and DBP using the weighted mode method (**Supplementary Table 7**). When the summary statistics of eGFRcys, another alternative kidney function marker, was used for the selection of genetic instruments for eGFRcr and BUN, we also observed similar significant causal estimates for eGFRcr and BUN on SBP and DBP using the weighted mode method (**Supplementary Table 8**).

### Selection of genetic instruments for SBP and DBP

Of the 551 reported index SNPs of SBP from UKB-ICBP, 494 remained after removing SNPs associated with potential confounders (**Supplementary Table 1, Figure 1).** After LD clumping and the checking of the effect alleles in the GWAS summary statistics of the exposure and outcome, 250 index SNPs remained. When eGFRcr was used as the outcome, Steiger filtering removed 10 index SNPs including those at *UMOD/PDILT* and *PRKAG2*, resulting in 240 genetic instruments for SBP (**Supplementary Table 3**). Of the reported DBP index SNPs (n = 537), 480 remained after removing SNPs associated with potential confounders and then 251 remained after LD clumping and checking of the effect alleles. When eGFRcr was used as the outcome, Steiger filtering removed 8 index SNPs including those at *UMOD/PDILT* and *PRKAG2*, resulting in 243 genetic instruments for DBP (**Supplementary Table 3)**. With the same SNP selection algorithms, 248 SBP and 238 DBP genetic instruments were selected when CKD was the outcome, and 243 SBP and 234 DBP genetic instruments were selected when BUN was the outcome **Supplementary Tables 3 and 5**).

### Causal effect estimates of blood pressure on kidney function

We observed that the causal estimates of blood pressure on kidney function were generally not significant using weighted mode, our primary method. The effect estimate for each SD higher SBP was −0.09 SD in log(eGFRcr) (95% CI: −0.18 to −0.002; p = 4.71×10^−2^, **Figure 3, Supplementary Table 6**). Similar non-significant causal estimates were observed for DBP on the three kidney function outcomes (eGFRcr, CKD, and BUN) using weighted mode as well as MRMix, the methods most robust to horizontal pleiotropy (**Supplementary Table 6**). In contrast, using IVW-FE, the causal estimates were significant across all blood pressure and kidney function traits, which might be due to horizontal pleiotropy^15^. For example, using the IVW-FE method, the causal estimate indicated a 27% higher odds ratio for CKD per each SD higher SBP (OR: 1.27, 95% CI: 1.17, 1.37, p = 2.01×10^−8^).

**Figure 3.**
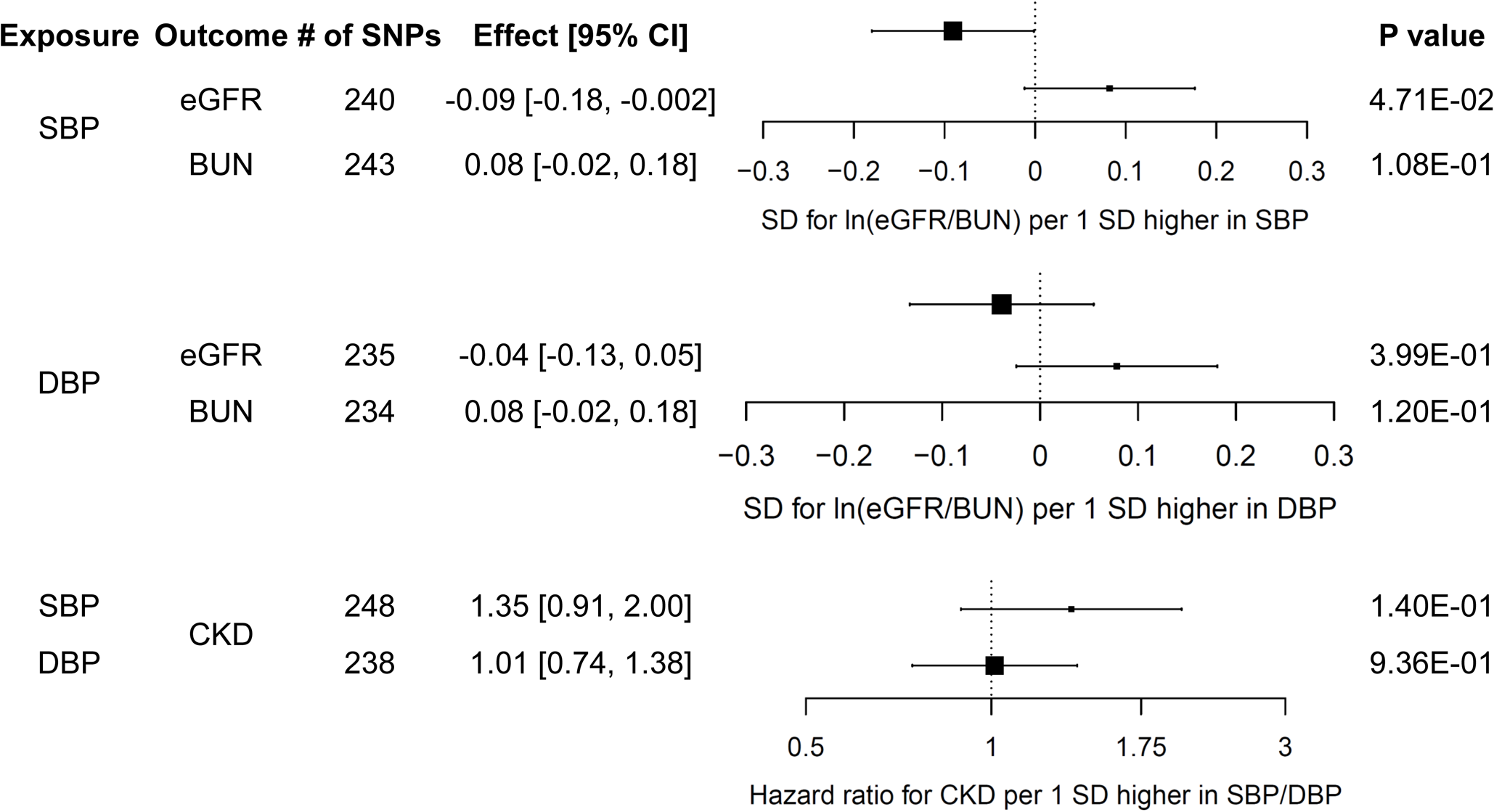
Estimates of the causal effects [95% confidence intervals] from SBP on eGFRcr and BUN (A), DBP on eGFRcr and BUN (B), and SBP and DBP on CKD (C) using the weighted mode method

Sensitivity analysis using blood pressure summary statistics from UKB only as exposure resulted in similar causal estimates for SBP and DBP to eGFR, BUN, and CKD (**Supplementary Table 7**). The scatter plots with the regression line from all Mendelian randomization analyses are presented in **Supplementary Figures 5a to 10a.** The forest plots of single SNP effects from blood pressure traits to kidney function traits are presented in **Supplementary Figures 5b to 10b**.

## DISCUSSION

Extensive Mendelian randomisation analyses, based on the largest GWAS summary statistics available to date on kidney function and blood pressure traits, showed evidence of a causal role of kidney function on blood pressure levels. Specifically, we observed that 50% lower eGFRcr results in 17.5 mmHg higher SBP and 8.4 mmHg higher DBP. In contrast, the causal role of blood pressure on kidney function levels were not supported across Mendelian randomisation methods. The significant causal effect of lower kidney function on higher blood pressure suggests preventing kidney function decline can reduce the public health burden of hypertension.

The finding of lower kidney function as causal to higher blood pressure is consistent with the genetics of hypertension-attributed kidney disease in African Americans, in whom the *APOL1* high-risk genotype confers twice the risk of CKD progression and appears to directly affect kidney function rather than blood pressure^47−49^. Few epidemiological studies reported lower kidney function as a risk factor for higher blood pressure^11 12^. Our finding of significant causal effects of lower kidney function on higher blood pressure suggests more research on the relation between kidney function and blood pressure before the development of either CKD or hypertension may increase our understanding on the interplay between these two diseases.

Using the Mendelian randomisation method, a study reported significant causal effects of higher albuminuria, an indicator of kidney damage, on higher SBP and DBP and vice versa^16^. These results are consistent with our findings of the significant causal effects of lower kidney function (eGFRcr and BUN) on higher SBP and DBP. In our study, causal effects of SBP and DBP on eGFR and BUN were inconsistent across Mendelian randomisation methods. Another Mendelian randomisation study reported significant causal effects of higher SBP and DBP to CKD using the IVW-FE method, which provides consistent estimates assuming that the sum of horizontal pleiotropic effects of all instruments is zero and horizontal pleiotropic effects are independent of instrument strength across all variants^50^. When applying the IVW-FE method in our study, the causal estimates of SBP and DBP on CKD were also significant and similar to those previously reported^15^. The weighted median method, which is known to be robust under the assumption that at least 50% of the selected instruments are valid, also showed some evidence of these causal effects. However, neither weighted mode nor MRMix, two methods which are known to be the most robust in the presence of complex pleiotropic effects, showed any evidence of statistical significance for these causal effects.

eGFRcr and BUN measures are challenging as exposure or outcome for Mendelian randomisation studies given that these measures have systematic measurement errors due to GFR biomarker-specific genetic determinants that are independent of kidney function, such as variants at *GATM* for creatinine metabolism^20 42 44^ and *SLC14A2* related to urea transport for BUN^45 46^. In two prospective observational studies that reported significant association between lower kidney function and incident hypertension, the kidney function biomarkers that significantly associated with hypertension were cystatin C and beta-2 microglobulin, whereas serum creatinine, the most commonly used GFR biomarker for kidney function estimation, was not significant^11 12^. In our Mendelian randomisation study, when eGFRcr or BUN measure were used as exposure, we used alternative kidney function biomarker to select genetic instruments that are more likely to reflect GFR. When blood pressure traits were used as the exposure, the systematic measurement errors of the kidney function traits due to GFR biomarkers may have biased the causal estimates to null.

Our use of Steiger filtering, which compared the effect size of a genetic instrument for exposure and outcome, suggested that some kidney function loci may affect kidney function through blood pressure, such as *FGF5*, and some blood pressure loci may affect blood pressure through kidney function, such as *UMOD*, which expresses exclusively in the kidney^51^. These results provide insight into the potential pleiotropy underlying GWAS findings of these traits.

Our study has several strengths. We used summary statistics from large-scale GWAS for evaluating causal effects. Using a power calculation method tailored for Mendelian randomisation, we have 90% power to detect a causal effect ≥ 0.013 SD difference in SBP per 1 SD difference in natural log-transformed eGFRcr and a causal effect ≥ 0.018 SD difference in natural log-transformed eGFRcr per 1 SD difference in SBP at an alpha level of 0.05. To overcome the systematic measurement errors due to the GFR biomarker components in eGFRcr and BUN measures, we used alternative GFR biomarkers to select genetic instruments that are more likely to reflect kidney function rather than biomarker metabolism. To reduce the possibility of violating the assumptions of Mendelian randomisation, we employed a range of techniques: evaluation of the association of index SNPs with potential confounders, use of Steiger filtering to reduce potential reverse causation driven by genetic instruments, and selecting a primary method that is robust to the presence of pleiotropy accompanied with sensitivity analysis with several alternative methods.

Some limitations warrant mentioning. In our primary analysis, the cohorts in CKDGen and UKB-ICBP had some overlap, which might lead to bias in the causal estimates^41^. However, our results using non-overlapping populations in exposures and outcomes (CKDGen and UKB only) were similar to our primary analysis. The CKDGen populations included cohorts of CKD patients and children. However, these cohorts only made up a small proportion of the European-ancestry study population. Overall, the populations in the summary statistics for exposures and outcomes were similar^52^.

In summary, using genetic instruments, we found that lower kidney function is causal to higher blood pressure. This result suggests that preventing kidney function decline may reduce the public health burden of hypertension.

## SUMMARY BOX

### What is already known on this topic

Lower kidney function has been associated with higher blood pressure and vice versa based on results from observational studies. It remains unclear whether these relations are causal.

### What this study adds

Higher kidney function has significant causal effects on lower blood pressure. These results suggest preventing kidney function decline can reduce the public health burden of hypertension.

## ACKNOWLEDGEMENT

The authors thank the Chronic Kidney Disease Genetics (CKDGen) Consortium (https://ckdgen.imbi.uni-freiburg.de), the UK Biobank (https://www.ukbiobank.ac.uk/), and the International Consortium of Blood Pressure-Genome Wide Association Studies (ICBP) for publicly sharing the genetic data we used in our causal analysis.

## FOOTNOTES

### Competing interests

All authors declare: no support from any organisation for the submitted work other than detailed above; no financial relationships with any organisations that might have an interest in the submitted work in the previous three years; no other relationships or activities that could appear to have influenced the submitted work.

### Funding

AK is supported by a Heisenberg Professorship (KO 3598/3-1, 3598/5-1) as well as CRC 1140 project number 246781735 and CRC 992 of the 10 German Research Foundation. NC is supported by National Human Genome Research Institute (NHGRI) grant R01 HG010480-01. AT is supported by National Institute of Arthritis and Musculoskeletal and Skin (NIAMS) grant R01AR073178-01A1. The Atherosclerosis Risk in Communities (ARIC) study is funded by HHSN268201100009C. The funding sources had no role in: a. the design or conduct of the study, b. the collection, management, analysis, and interpretation of the data, or c. preparation, review, or approval of the manuscript.

### Contributors

YZ, JC, NC, and AT designed the study, wrote the research plan, and interpreted the results. YZ and AT wrote the first draft of the manuscript with critical comments and revision from JC, GQ, MEG, EB, HS, AT, CP, AK, and NC. AT is the guarantor. The corresponding author attests that all listed authors meet authorship criteria and that no others meeting the criteria have been omitted.

### Ethical approval

No ethics approval was acquired for the analysis using publicly available data.

### Data sharing

All of the summary level data used are available for instant download at the public repositories. The statistical code is available from the first author and corresponding author at.

### Patient and Public Involvement statement

Patients and the public were not involved in this study.

## Notes

https://ckdgen.imbi.uni-freiburg.de/

http://www.nealelab.is/uk-biobank

http://eagle-i.itmat.upenn.edu/sweet/provider?uri=http://eagle-i.itmat.upenn.edu/i/00000155-dc32-22ed-627c-14a480000000

