## Supplementary Figures for "Kidney Function and Blood Pressure: A Bi-directional Mendelian Randomisation Study"

#### Supplementary Figure 1A. Regression lines of MR tests from eGFRcr on SBP

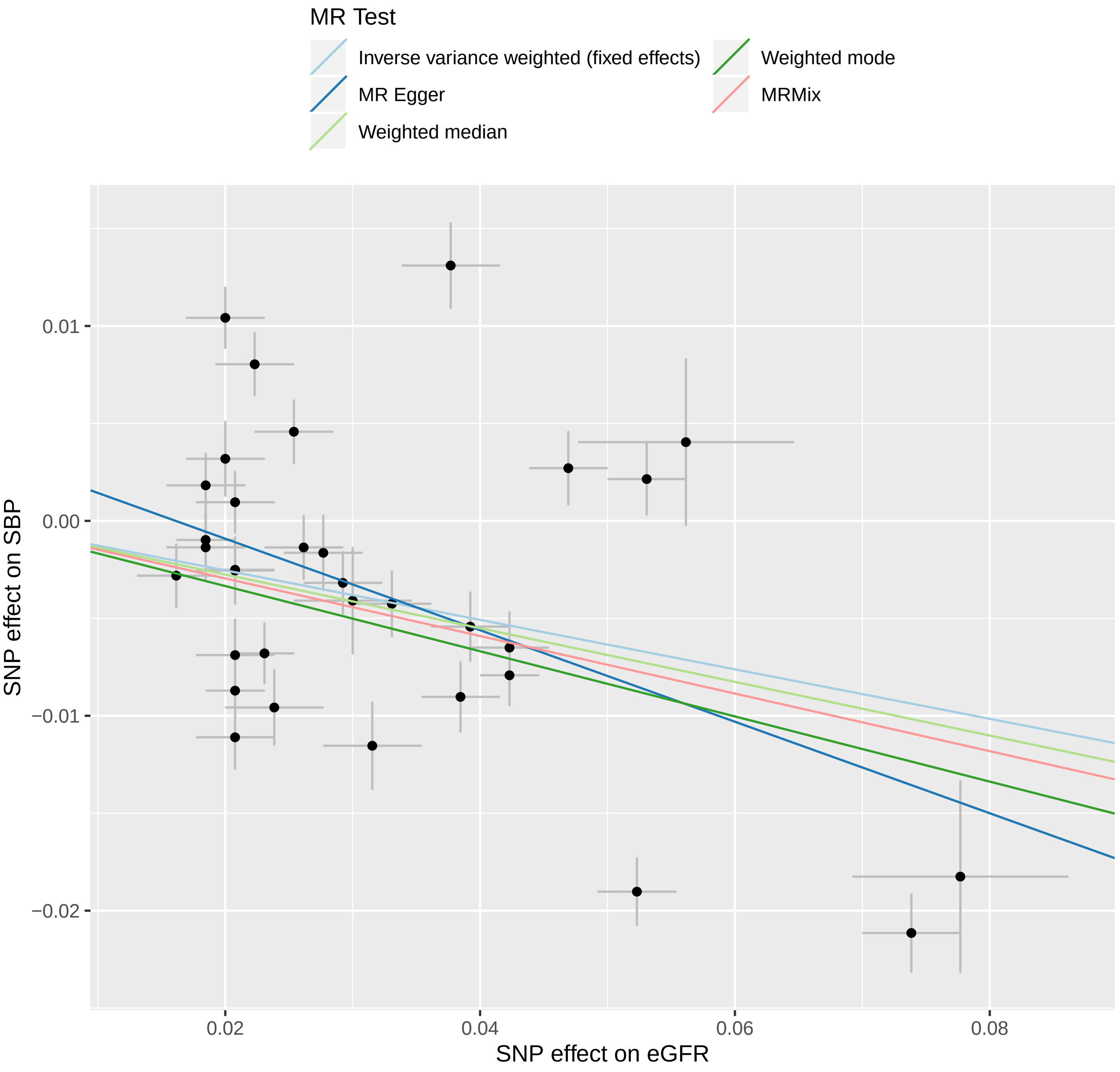

### Supplementary Figure 1B. Forrest plot of single SNP from eGFRcr on SBP

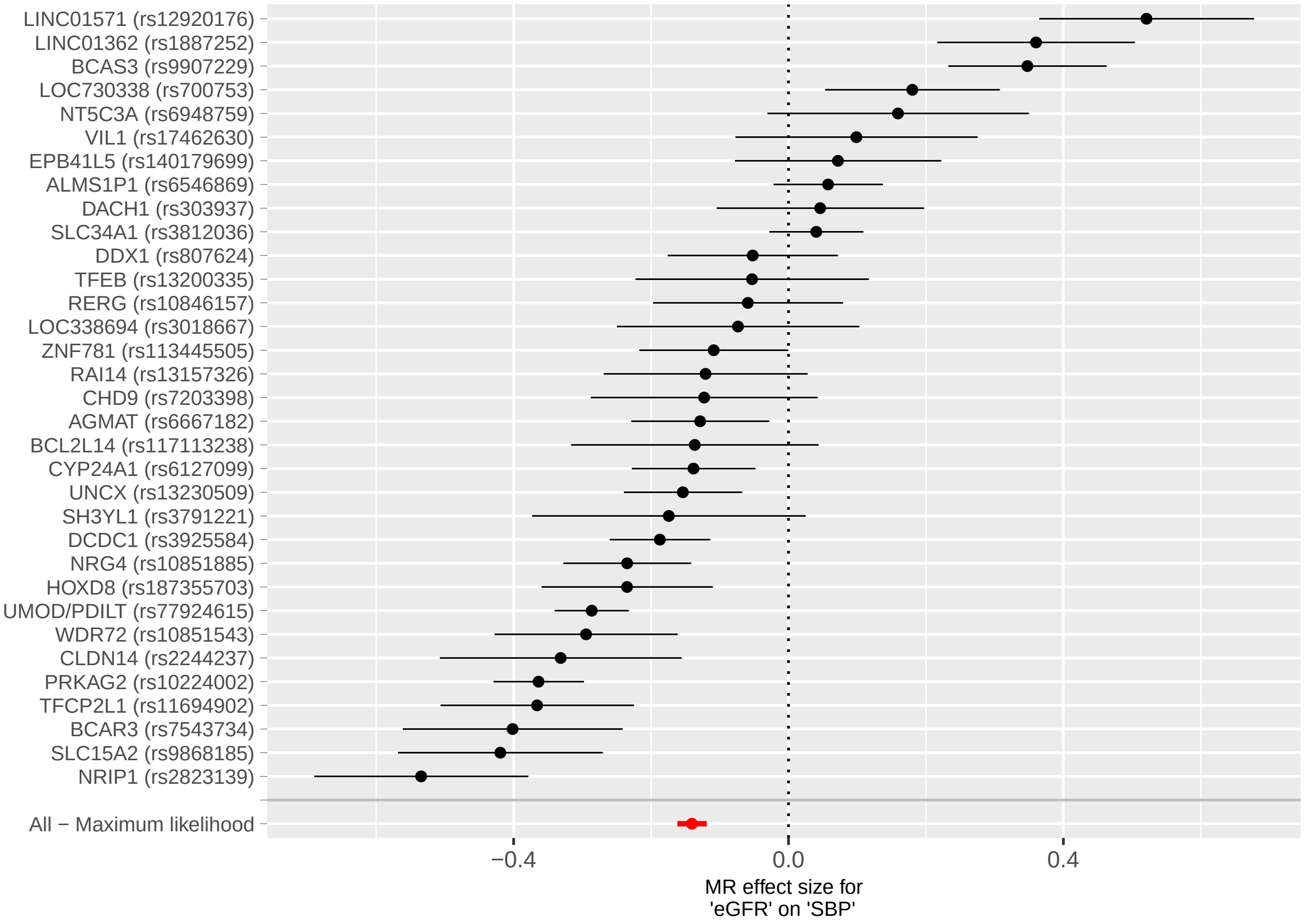

**Supplementary Figure 2A. Regression lines of MR tests from eGFRcr on DBP**

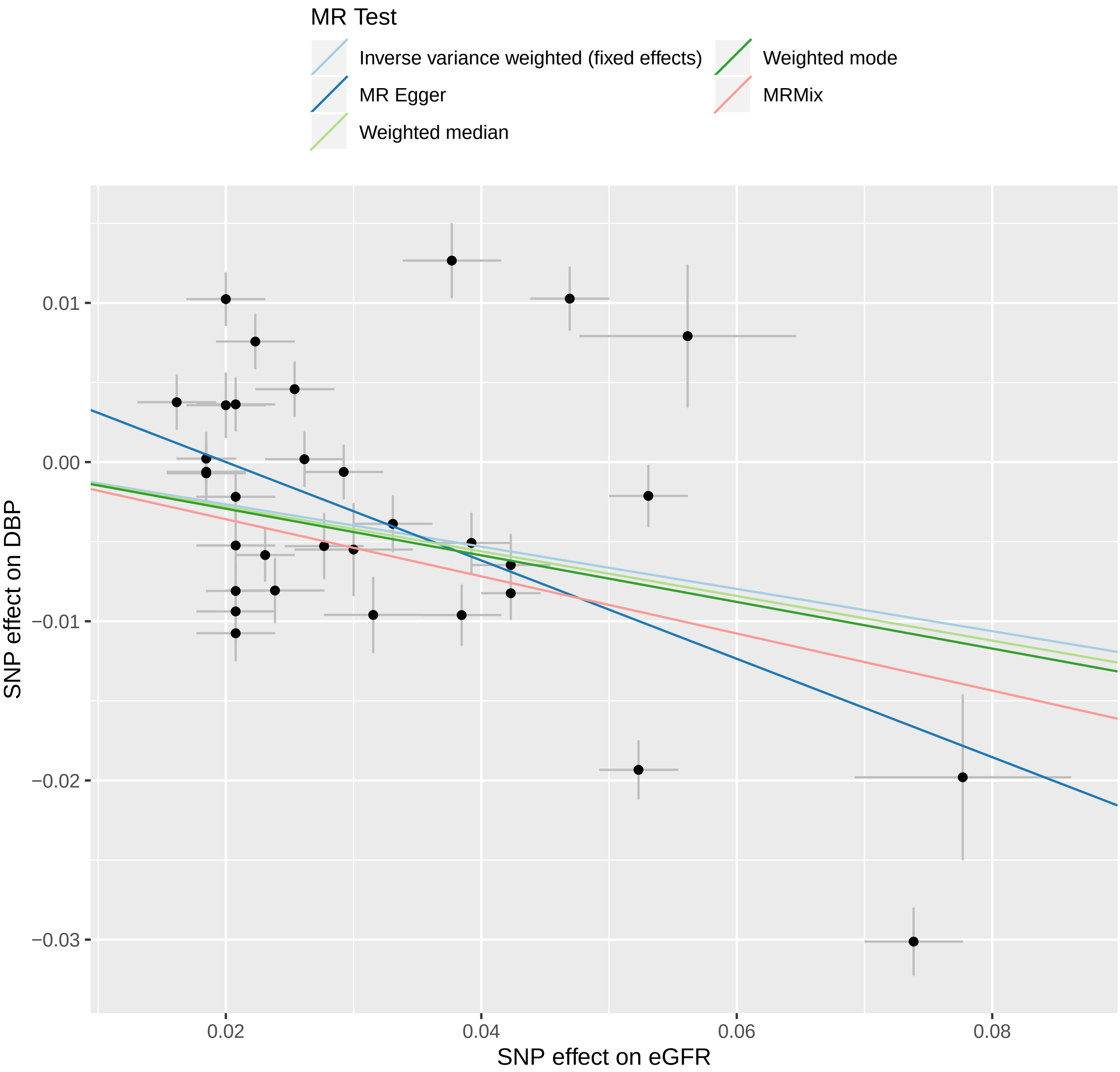

### Supplementary Figure 2B. Forrest plot of single SNP from eGFRcr on DBP

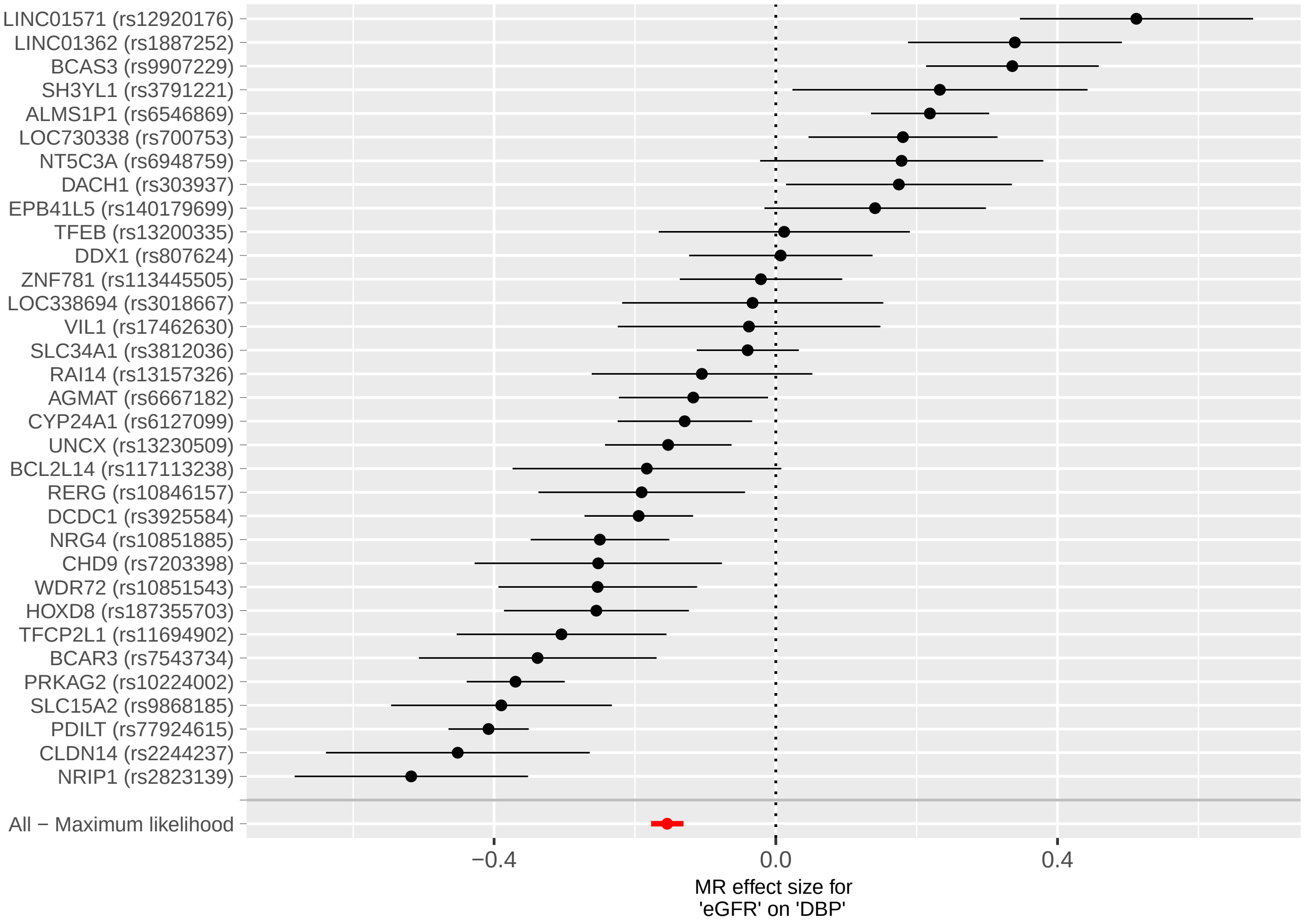

**Supplementary Figure 3A. Regression lines of MR tests from BUN on SBP**

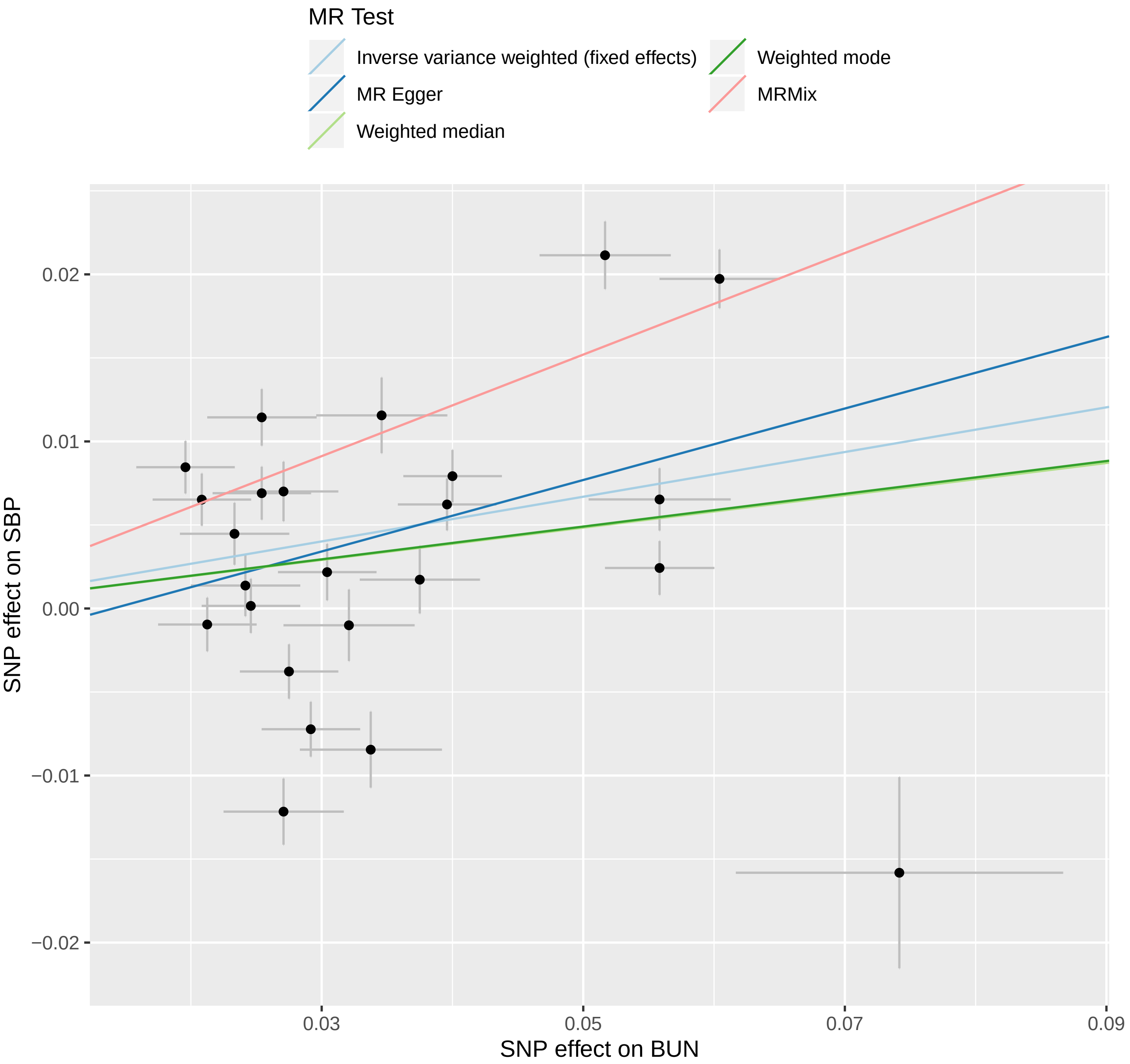

### Supplementary Figure 3B. Forrest plot of single SNP from BUN on SBP

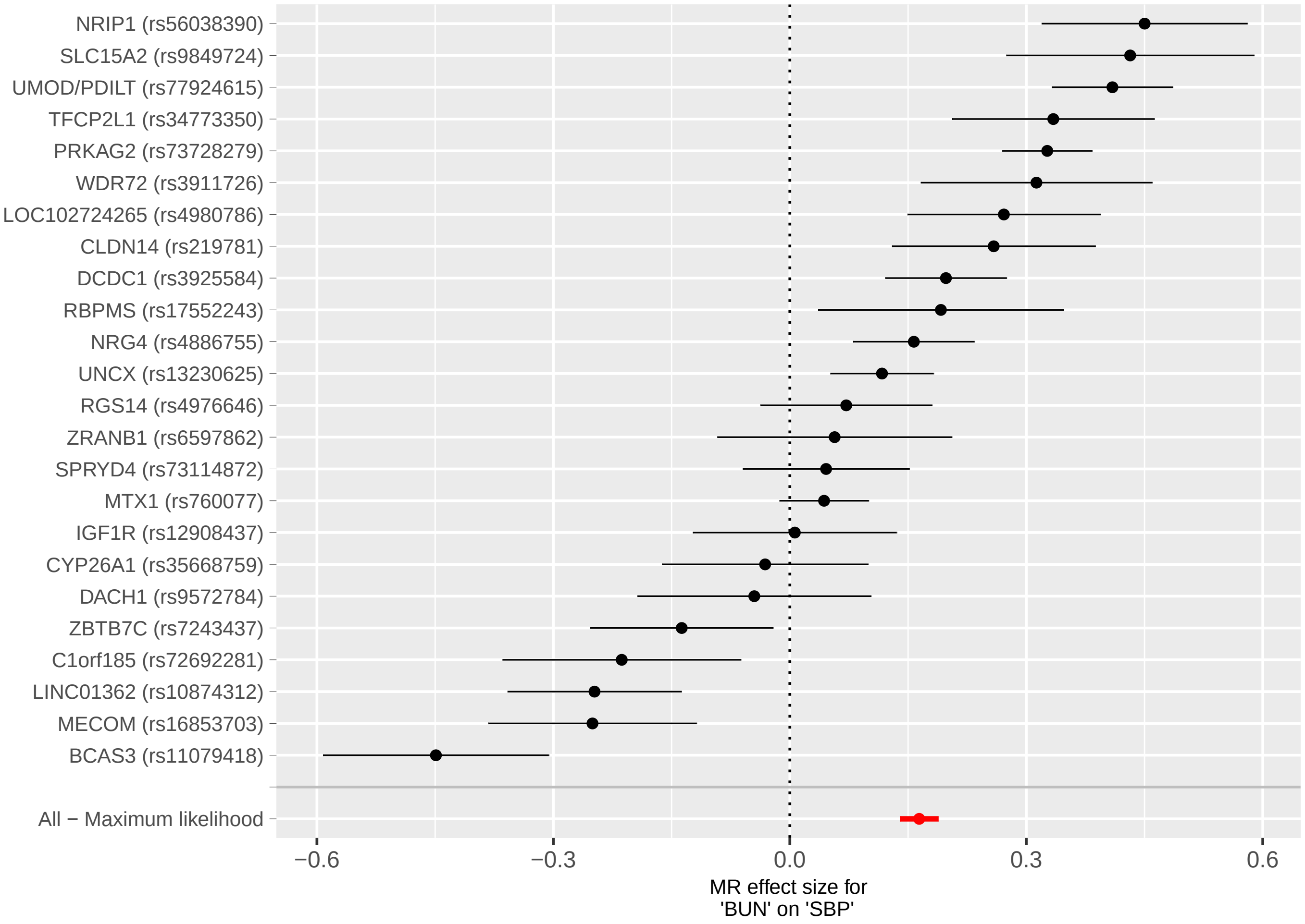

**Supplementary Figure 4A. Regression lines of MR tests from BUN on DBP**

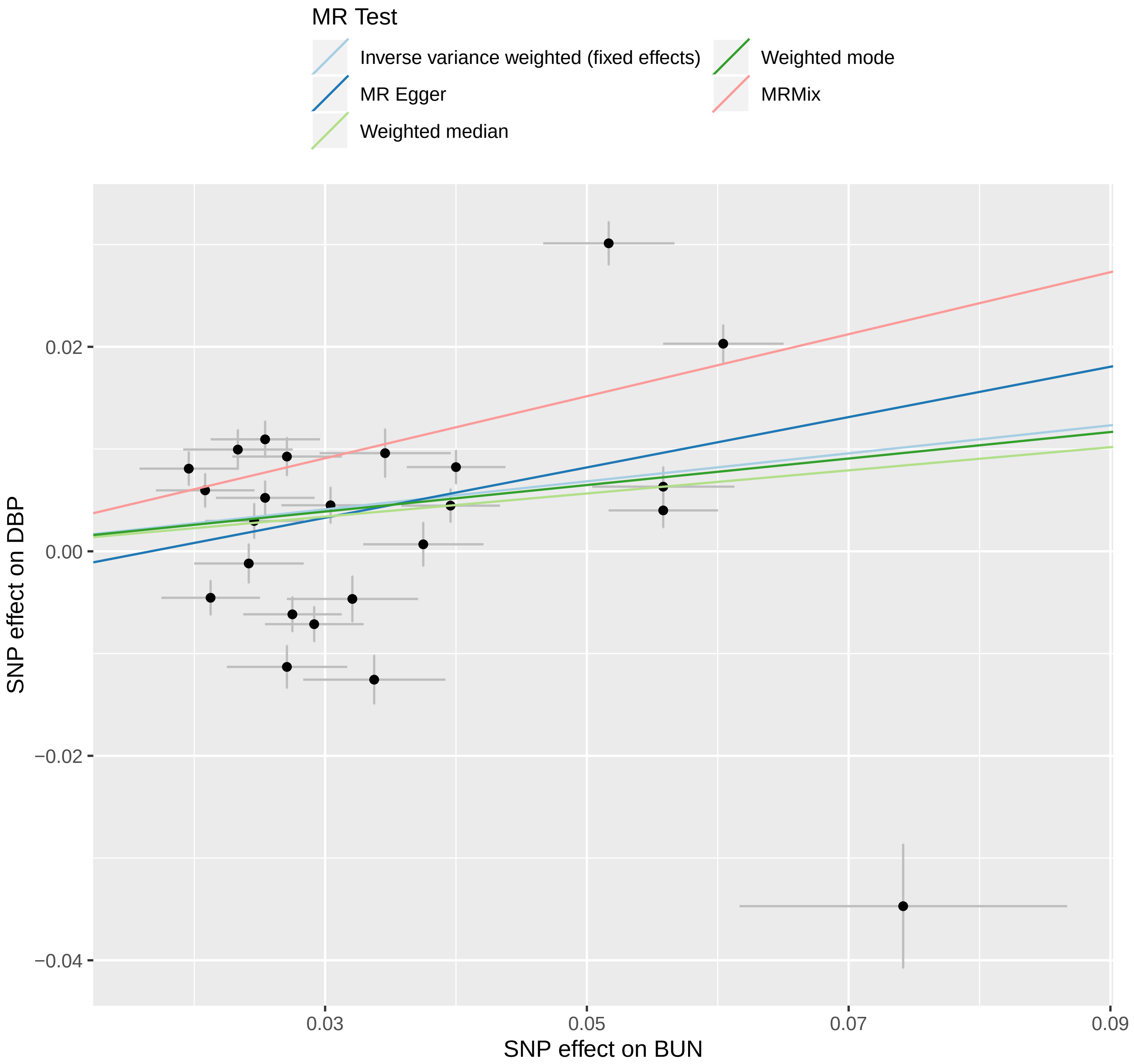

#### Supplementary Figure 4B. Forrest plot of single SNP from BUN on DBP

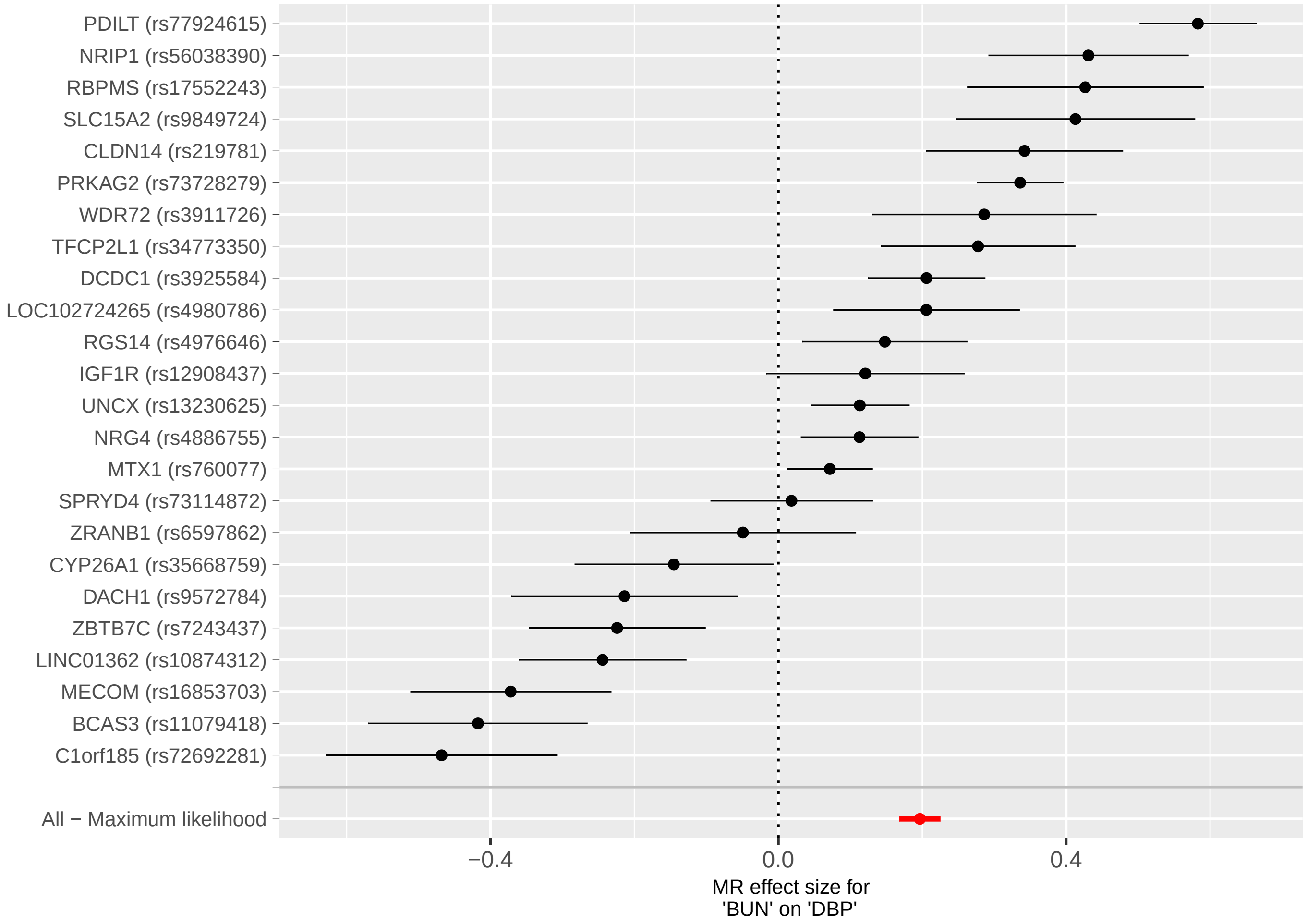

#### Supplementary Figure 5A. Regression lines of MR tests from SBP on eGFRcr

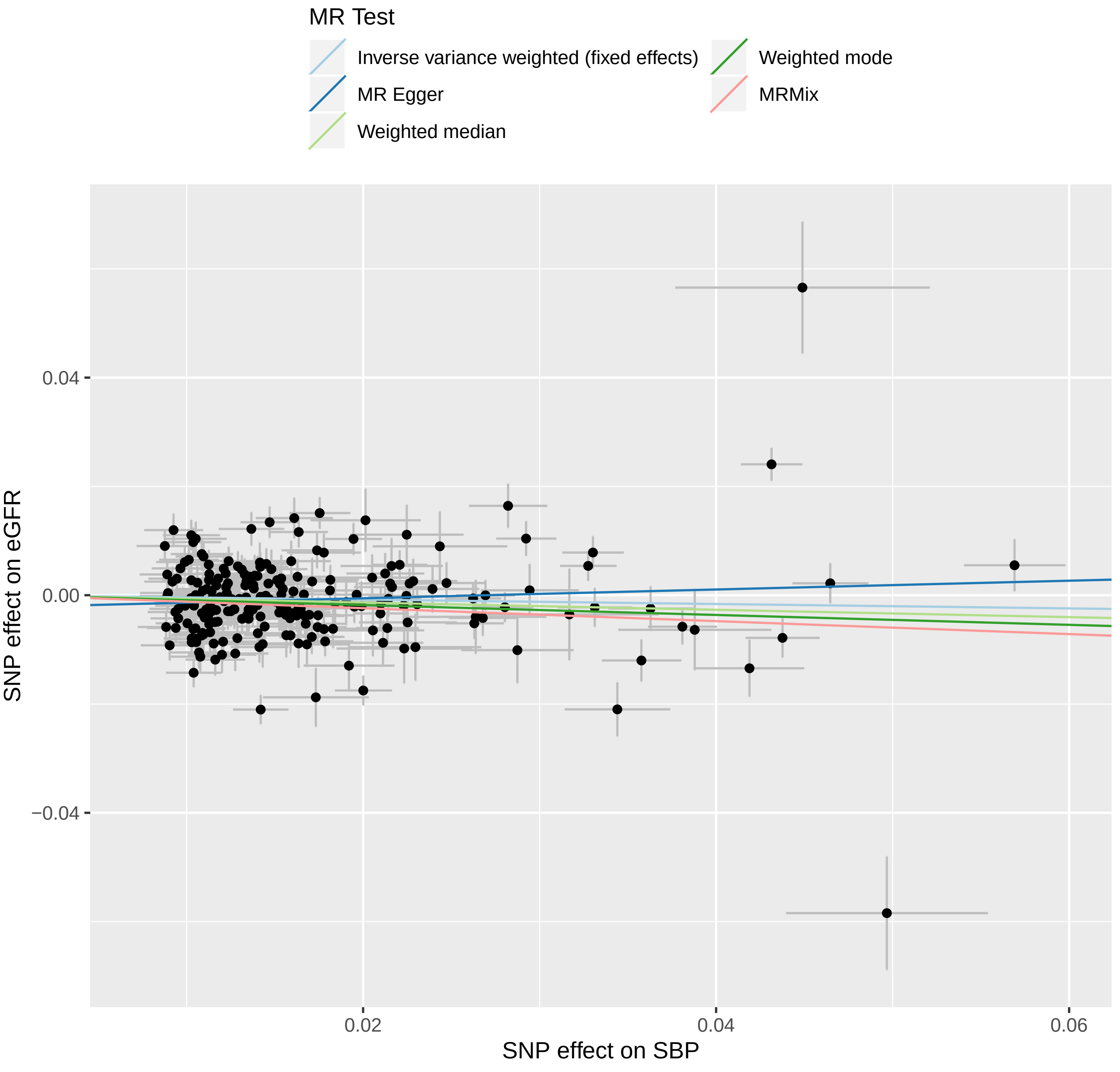

#### Supplementary Figure 5B. Forrest plot of single SNP from SBP on eGFRcr

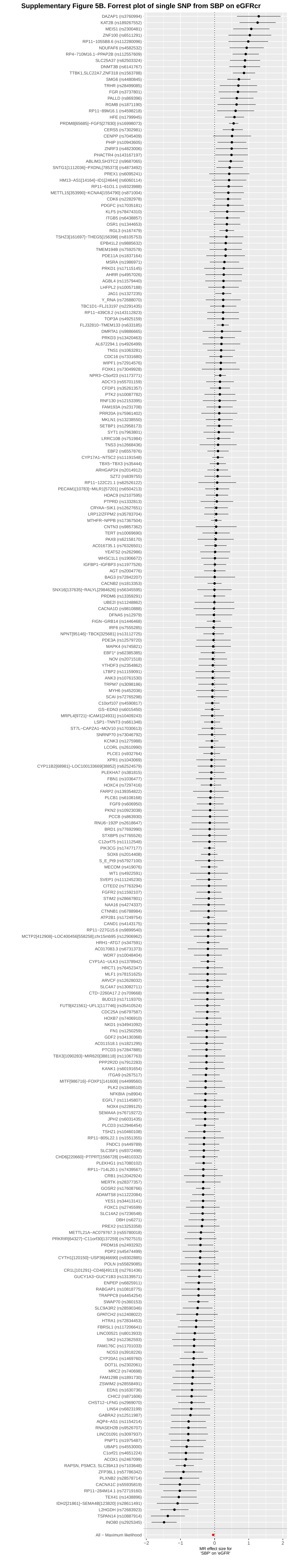

#### Supplementary Figure 6A. Regression lines of MR tests from SBP on CKD

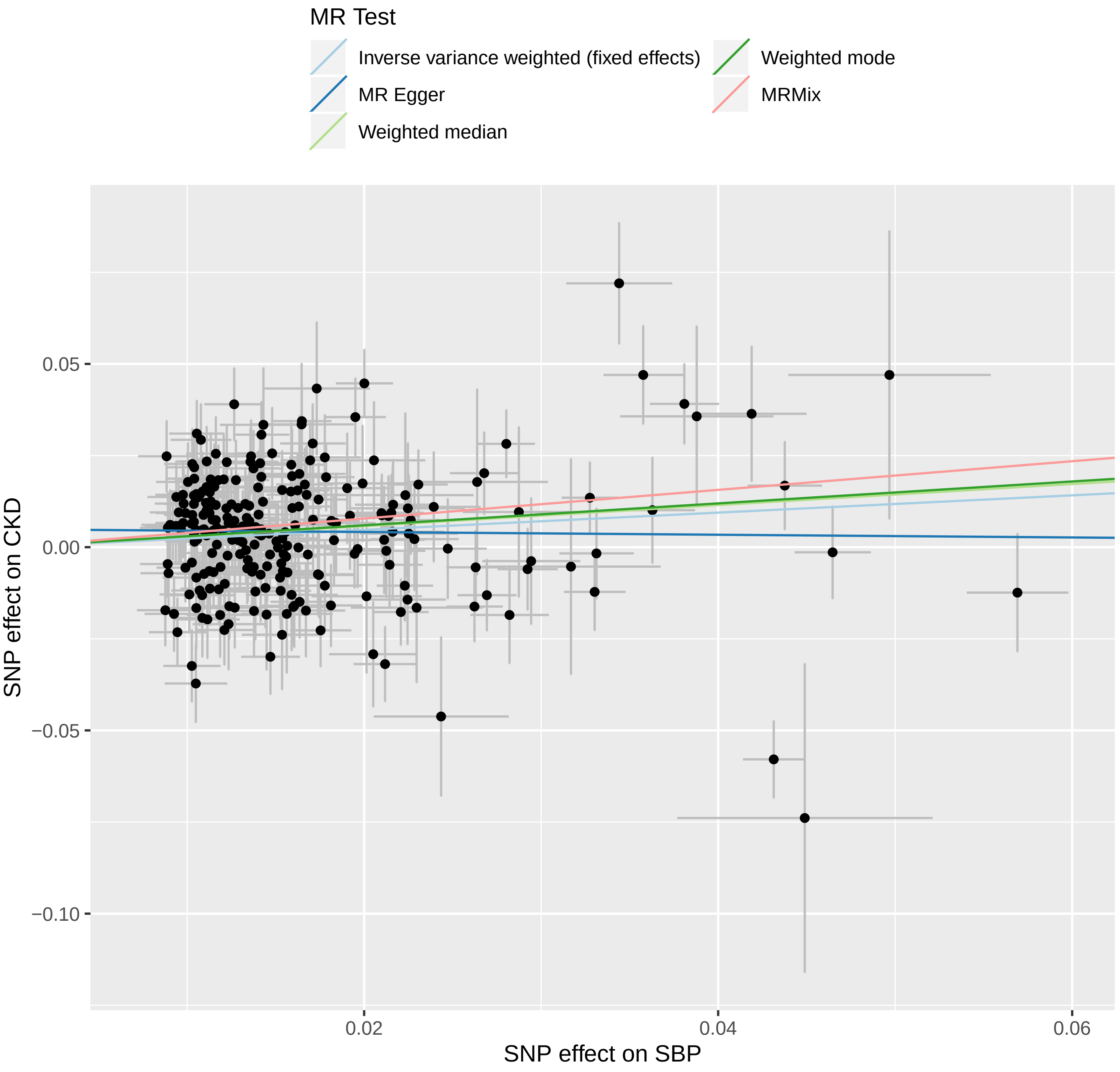

#### Supplementary Figure 6B. Forrest plot of single SNP from SBP on CKD

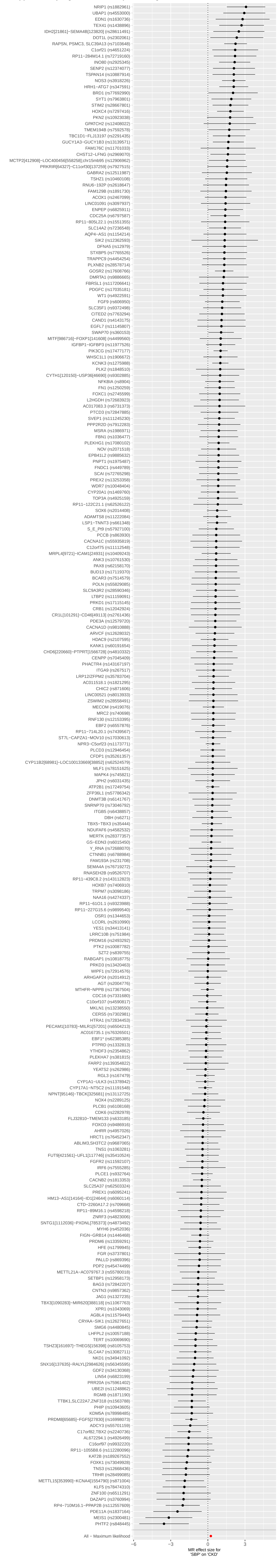

**Supplementary Figure 7A. Regression lines of MR tests from SBP on BUN**

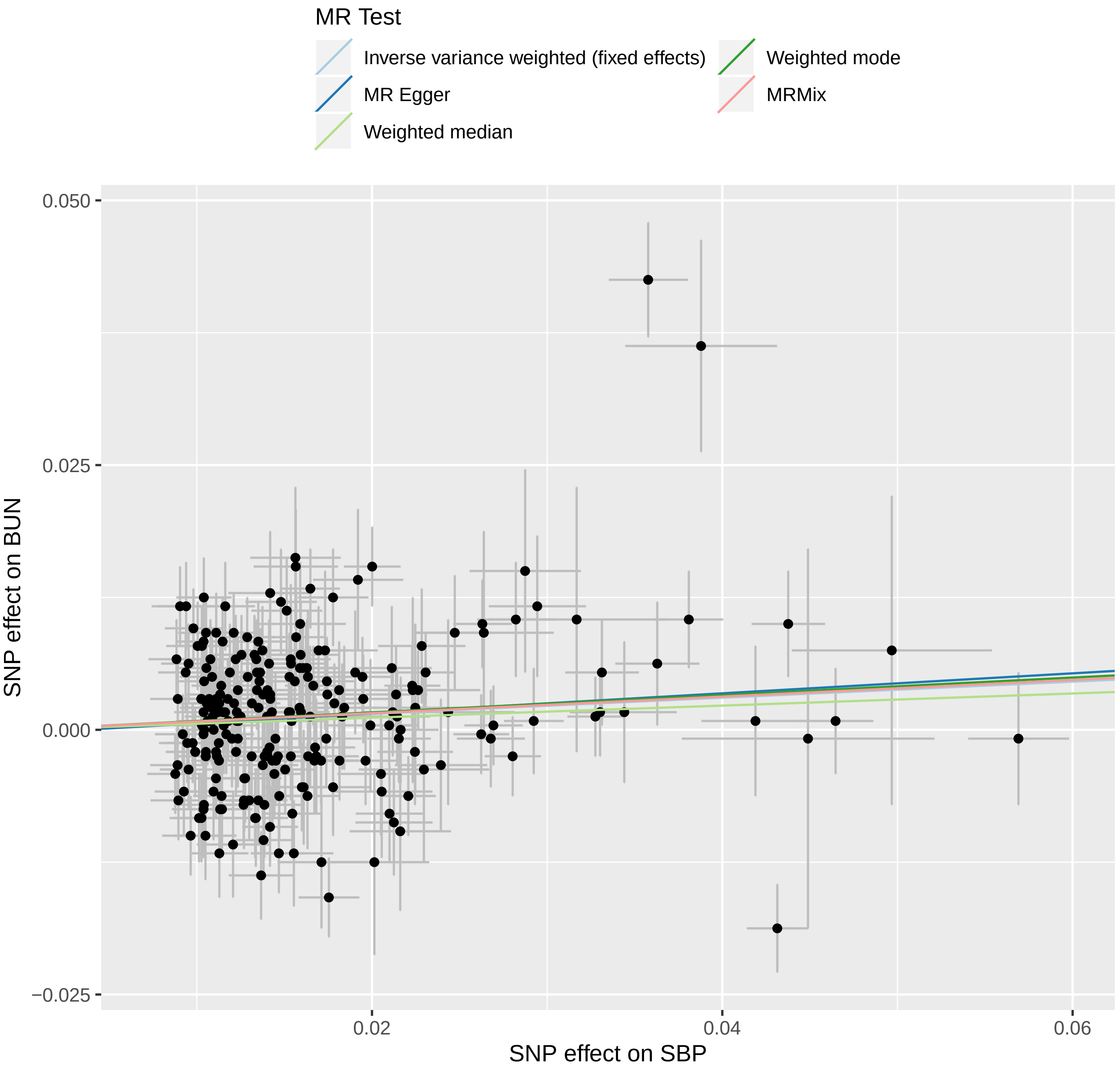

**Supplementary Figure 7B. Forrest plot of single SNP from SBP on BUN**

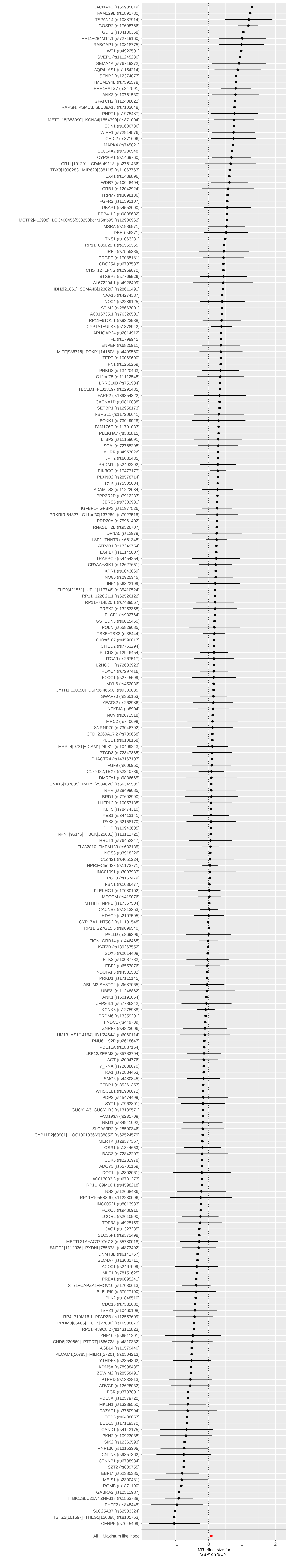

**Supplementary Figure 8A. Regression lines of MR tests from DBP on eGFRcr**

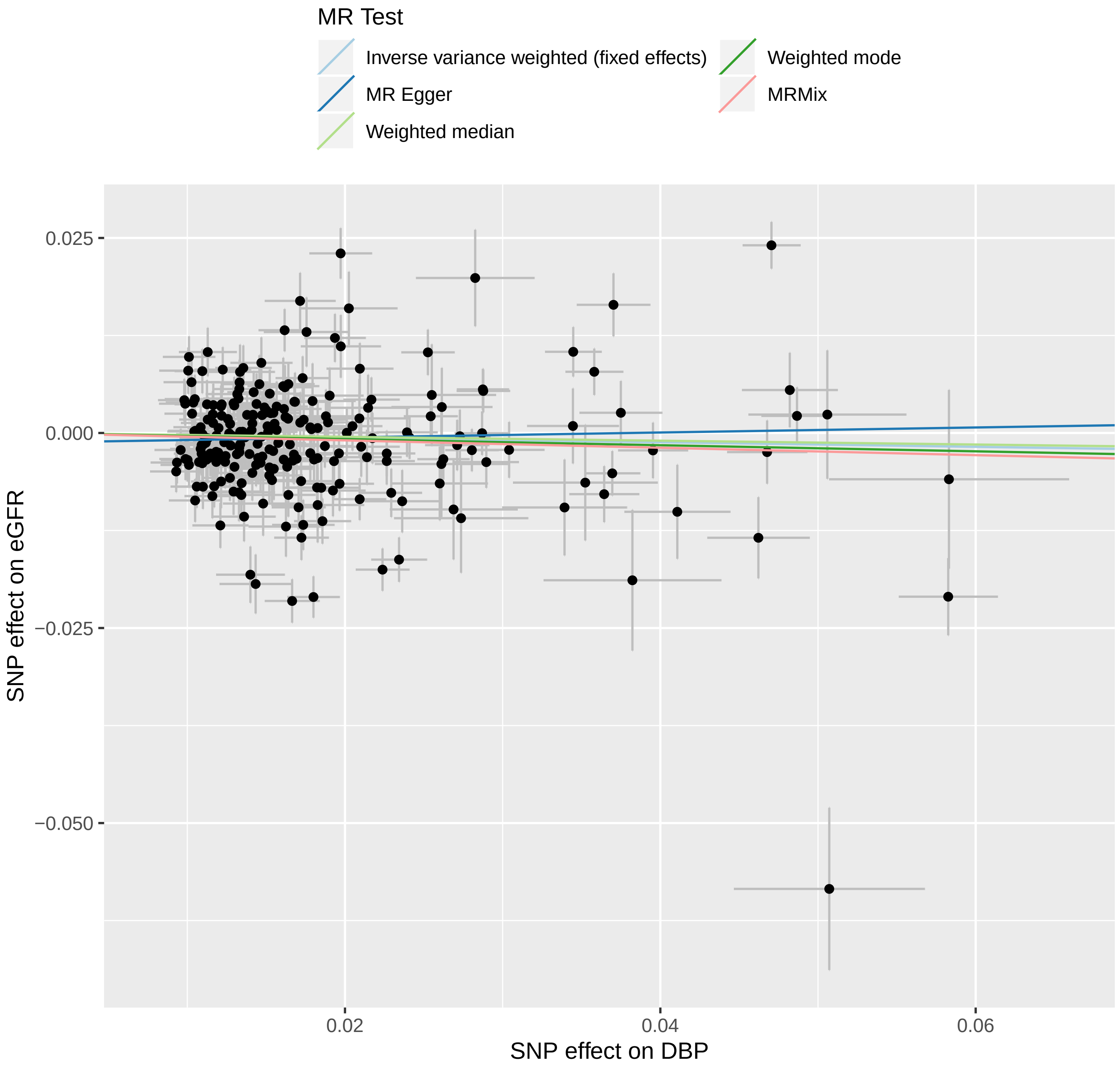

**Supplementary Figure 8B. Forrest plot of single SNP from DBP on eGFRcr**

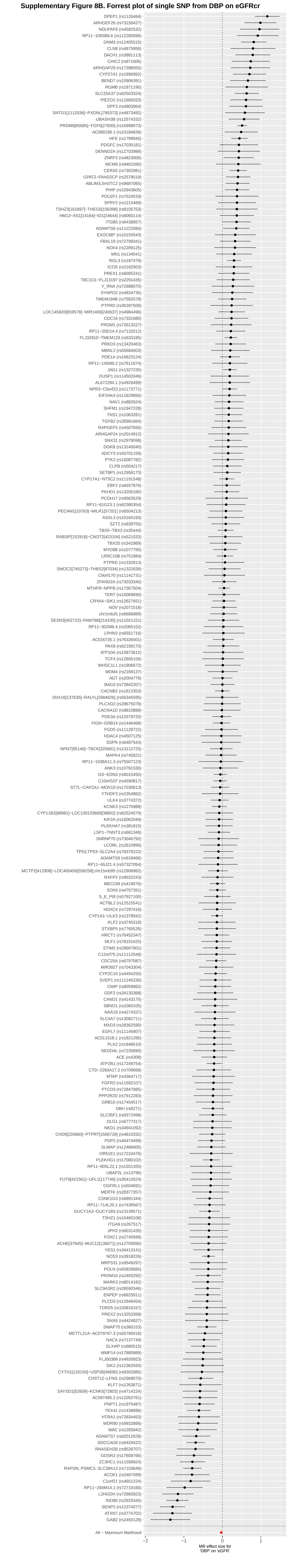

### Supplementary Figure 9A. Regression lines of MR tests from DBP on CKD

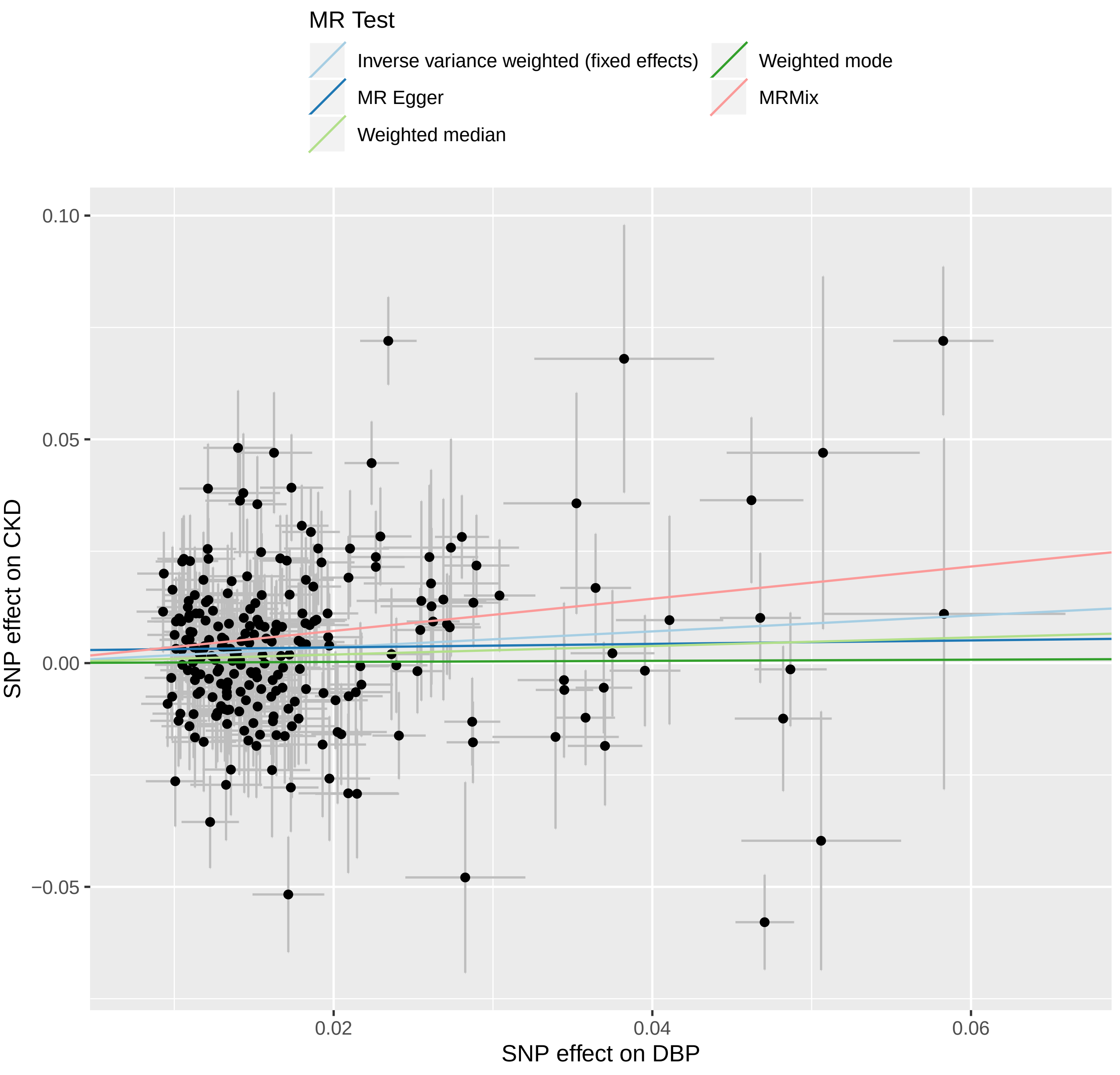

#### Supplementary Figure 9B. Forrest plot of single SNP from DBP on CKD

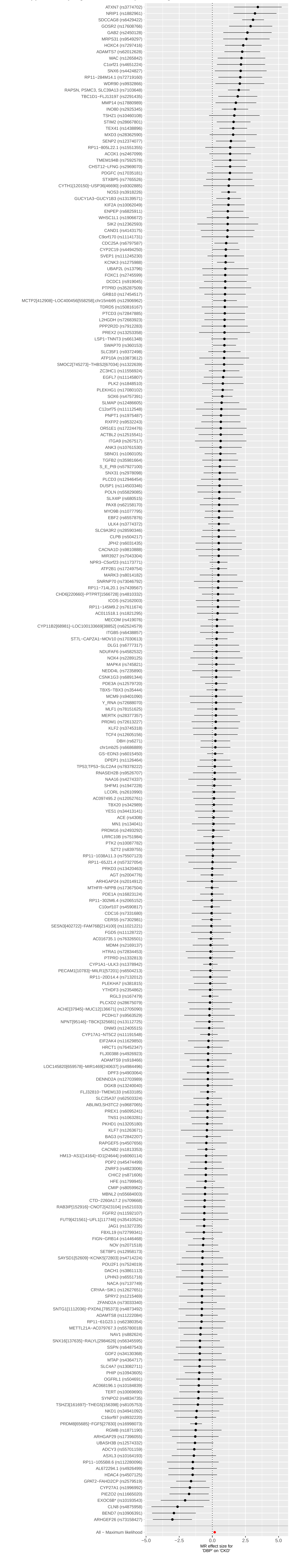

#### Supplementary Figure 10A. Regression lines of MR tests from DBP on BUN

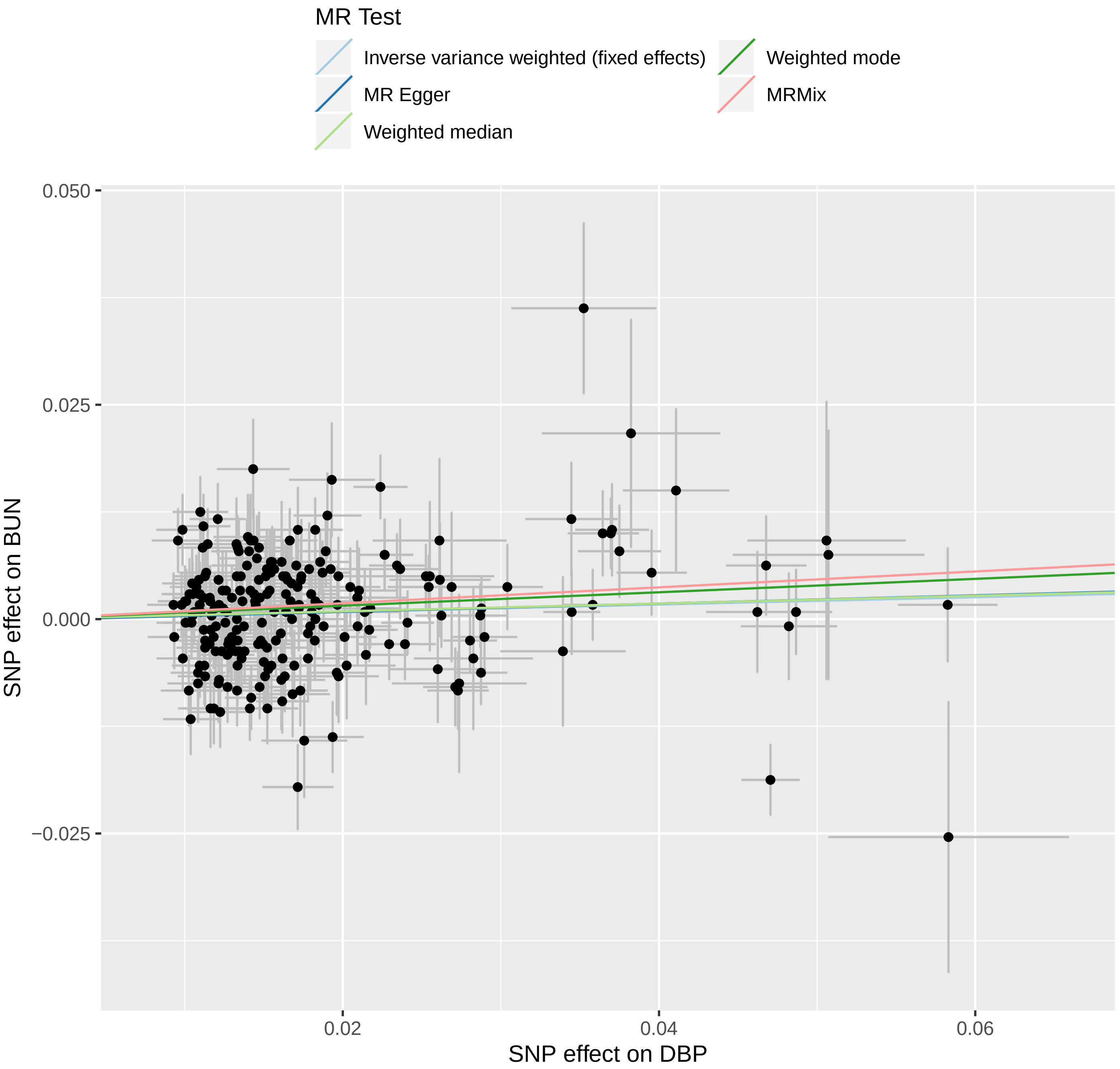

#### Supplementary Figure 10B. Forrest plot of single SNP from DBP on BUN

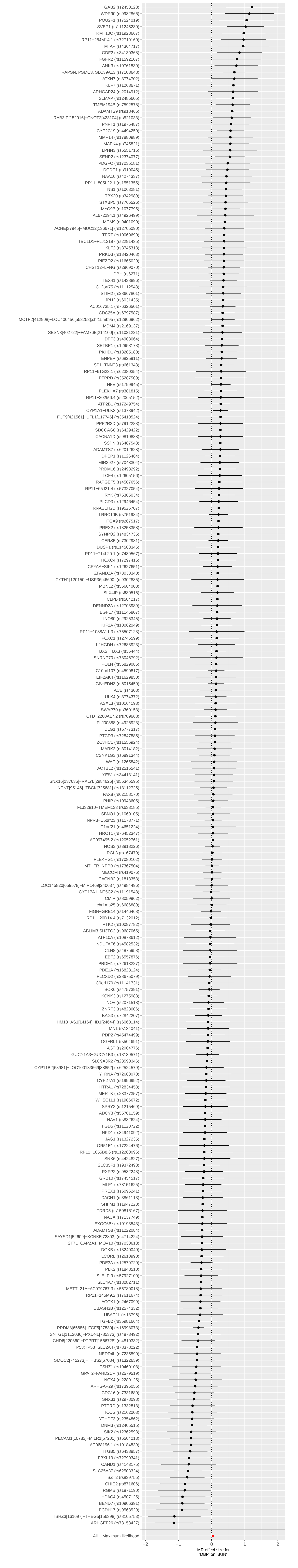
